# Auditory corticostriatal connections in the human brain

**DOI:** 10.1101/2022.08.04.502679

**Authors:** Kevin R. Sitek, Leah B. Helou, Bharath Chandrasekaran

## Abstract

Auditory learning depends on sensory, perceptual, decisional, and reward-based processes that are supported by the dorsal striatum. Auditory corticostriatal connections have been well-characterized in animal models including non-human primates, where primary auditory cortex preferentially connects to putamen, and caudate head receives most of its inputs from anterior superior temporal cortex. However, the extent to which human auditory corticostriatal connectivity follows similar organizational principles is challenging to assess due to the density of these striatal structures relative to the resolution of traditional diffusion MRI techniques, as well as dorsal striatum’s location near multiple major crossing white matter bundles. We leveraged high-quality diffusion-weighted MRI tractography to ‘virtually’ dissect structural pathways between auditory cortical regions and dorsal striatal regions in a sub-millimeter resolution single-participant dataset. Across most of auditory cortex, putamen connections were more frequent than caudate connections; only anterior-most superior temporal cortex had strong connectivity with caudate, specifically the caudate head. Putamen streamline endpoints were largely along the ventral portion of the structure, ranging from caudal to middle putamen. These results were consistent across analysis and tractography pipelines. In contrast to the auditory findings, visual corticostriatal streamlines did not preferentially reach putamen. We replicate these results in an independent sample of near-millimeter resolution single-session diffusion MRI from the Human Connectome Project. Overall, our results suggest strong structural connectivity between primary and association auditory cortices with putamen but not with any subdivision of caudate. Prioritized connectivity between superior temporal cortex and putamen is highly suggestive of distinct functional roles for striatal subdivisions in auditory perception.

## Introduction

Dorsal striatum is extensively involved in mediating learning and decision-making across motor, auditory, visual and cognitive domains (Groenewegen 2003; Lim et al. 2014; Ell 2011; Seger and Cincotta 2005; Seger 2013; Brovelli et al. 2011). Within the auditory system, corticostriatal pathways are argued to be key intermediaries involved in perceptual learning, categorization, and decision making (Znamenskiy and Zador 2013; Xiong et al. 2015; Lee et al. 2015). Corticostriatal pathways are hypothesized to underlie human auditory processes such as speech learning in adults (Chandrasekaran et al. 2014; Feng et al. 2019; Feng et al. 2021), language more broadly (Ullman 2001), and music (Zatorre and Salimpoor 2013).

Despite more recent interest in auditory corticostriatal function, very little is known about the anatomical pathways that connect the human auditory system to the striatal sub-structures. Existing information about auditory striatal circuits largely comes from non-human animal models (Yeterian and Pandya, 1998; Znamenskiy and Zador, 2013; Ponvert and Jaramillo 2019). Rodents are frequently used as model systems in neuroscience research (Hintiryan et al. 2016; Hunnicutt et al. 2016), including mapping of direct connections between auditory cortex and dorsal striatum (Ghosh and Zador 2021; Xiong et al. 2015; Znamenskiy and Zador 2013; Lee et al. 2015). However, neuroanatomical differences between rodents and humans make it challenging to translate corticostriatal connectivity patterns from rodents to humans; in contrast, humans are thought to retain much of the neuroanatomy of our non-human primate relatives, including corticostriatal organization (Balsters et al. 2020). Yet even between humans and macaques, differences in intrinsic organization of dorsal striatum are clearly documented (Liu et al. 2021).

But despite the likelihood of species-specific striatal connectivity patterns, to our knowledge, no focused mapping of human corticostriatal connectivity exists in the literature which can be compared to the extensive non-human primate literature.

To the extent that macaque and human striatum share similarities in organization relative to rodent models, we can make some predictions about the connectivity of the superior temporal cortical regions with both caudate and the putamen based on early neuroanatomical tracing literature in macaques (Yeterian and Pandya 1998; Yeterian and Van Hoesen 1978). In a series of experiments, investigators used anterograde autoradiographic tracers to map the corticostriatal pathways. In particular, temporal neocortical regions that are cortico-cortically connected with frontal cortex were also preferentially linked to the caudal portions of the putamen, above and beyond what is seen in other regions of interest in the frontal, parietal, and occipital lobes (Yeterian and Van Hoesen 1978). Later research took a more finely parcellated approach by distinguishing between anatomical regions within the superior temporal plane versus the physiologically-defined regions of primary and association auditory cortices (Galaburda and Pandya 1983). These studies verified prior findings (Kemp and Powell, 1970) that primary auditory cortex (AI) has limited projections to the dorsomedial portion of the tail of the caudate and the adjacent caudoventral segment of the putamen (Yeterian and Pandya 1998). Injections of tracer into associative auditory cortex (AII) revealed substantially more widespread corticostriatal projections that were distributed along the ventral aspect of the body of the caudate, the medial sector of its tail, and the putamen both rostro-ventrally and caudo-ventrally. With placement of tracer caudal to AI in the supratemporal plane, striatal labeling occurred in the head and body of the caudate and in rostral and caudal aspects of the putamen. Labeling patterns differed from the superior temporal gyrus compared to these regions in the supratemporal plane. Of all the regions studied, AI showed the most modest corticostriatal connectivity and the belt areas showed more robust connections. This pattern was similar to what was shown in neuroanatomical studies of primary visual corticostriatal connections (Kemp and Powell, 1970), supporting the conclusion that the functions of primary auditory cortex might have limited dependence on the striatum as compared to the belt regions and superior temporal gyrus. The tail of the caudate nucleus and the caudal putamen occupied the most substantial projection zone of the superior temporal cortex, implicating corticostriatal connections in the functionality of these regions, e.g., sound recognition, encoding vocalizations of conspecifics, sensorimotor association, and localization of sound sources. More recent work has described this “tail of the striatum” as a unique subdivision of dorsal striatum with an integrative role in sensory processing (Valjent and Gangarossa 2021; Cox and Witten 2019).

Direct connections from sensory cortex to dorsal striatum is also evident within the somatosensory system, where connections to putamen were traced from specific somatosensory regions in squirrel monkeys (Flaherty and Graybiel 1994; Parthasarathy and Graybiel 1997). Outputs from these somatosensory-receiving putamen regions converged in globus pallidus, suggesting a sensorimotor convergent zone in striatum. Flaherty and Graybiel proposed that one function of striatal modularity is to split input from one functional cortical region and thus initiate a temporary divergence—a striatal association network. In the auditory system, such a network would likely be built upon the widespread connections from associative auditory cortex throughout dorsal striatum discussed in the previous paragraph (Yeterian and Pandya 1998).

While divergent striatal association connections have not been clearly delineated in the auditory system, there is strong evidence for separate pathways with distinct functionality within auditory cortex, where dual pathways corresponding to “what” (ventral) and “where” (dorsal) processing were originally identified in primates (Romanski et al. 1999; Rauschecker and Tian 2000; Rauschecker and Scott 2009; Kaas and Hackett 2000). A recent fMRI study in macaques is more suggestive of auditory learning-dependent activity within putamen that perhaps takes advantage of sensorimotor convergent zones (Archakov et al. 2020). This study found greater putamen (as well as motor cortical) activation when the animals listened to sound patterns they had learned to produce compared to sound patterns that were unfamiliar to them.

In contrast to the methods available in non-human primates and other animal models, investigating human corticostriatal connectivity predominantly necessitates non-invasive approaches, particularly diffusion MRI tractography. Diffusion MRI is sensitive to the motion of water molecules in the brain, which is constrained by white matter and thus allows us to infer the orientation of white matter within each diffusion MRI voxel. We can then traverse from voxel to voxel to estimate pathways of white matter connectivity, an approach known as tractography. While diffusion MRI in general and tractography in particular have made tremendous progress over the past two decades, including auditory thalamocortical connections via the acoustic radiations (Maffei et al. 2019a; Maffei et al. 2018; Maffei et al. 2019b), investigations into human corticostriatal connectivity have been more limited (Ford et al. 2013; Avecillas-Chasin et al. 2016; Marrakchi-Kacem et al. 2013; Calabrese et al. 2022; Zhang et al. 2017; Waugh et al. 2022; Feng et al. 2019; Ghaziri et al. 2018). The rich interconnectedness of striatal subregions provides a challenge to diffusion MRI data with lower spatial and angular resolution, as well as to tractography methods that estimate only a single white matter orientation (such as diffusion tensor imaging, or DTI). Further, studies that do touch upon connectivity between superior temporal cortex (where auditory cortex resides) and dorsal striatum tend to focus on specific superior temporal sub-regions (Waugh et al. 2022; Feng et al. 2019), as opposed to fine-grained parcellations of all of superior temporal cortex similar to the macaque literature (Yeterian and Pandya 1998); striatal segmentations are also typically coarse-grained (although see (Tian et al. 2020) for a recent gradient-based approach to striatal sub-segmentation). Thus, with growing recognition of the importance of corticostriatal connections for human audition (and more broadly, communication)—and their potential dysfunction in autism (Abrams et al. 2016; Di Martino et al. 2011), developmental language disorders (Krishnan et al. 2016), persistent developmental stuttering (Giraud et al. 2008; Alm 2004; Chang and Zhu 2013), and Parkinson’s Disease (Jafari et al. 2020)—we need to better understand fundamental connectivity patterns between superior temporal cortex and dorsal striatum (Yi et al. 2019).

In the current study, we leveraged high resolution diffusion MRI tractography to virtually dissect auditory cortical–striatal connections in human participants. In particular, we analyzed one of the highest quality in vivo human datasets publicly available, collected over eighteen hours in a single individual on the diffusion MRI-optimized Siemens Connectom scanner with over 2800 diffusion directions and spatial resolution below 1 mm isotropic. We first mapped connections between subdivisions of auditory cortex in superior temporal cortex with the principal dorsal striatal input regions: caudate and putamen. We considered both tractography endpoints as well as white matter pathways of corticostriatal networks. Based on animal literature (Yeterian and Pandya 1998), we predicted strong tractography connectivity between primary and secondary auditory cortices in posterior superior temporal cortex and caudal striatal subdivisions (posterior putamen and caudate tail), while increasingly associative anterior superior temporal regions would be better connected with rostral striatum (anterior putamen and caudate head). Next, to explore the possibility of parallel organization across sensory modalities, we compared corticostriatal connectivity in the auditory system with homologous visual corticostriatal connectivity. Previous research in animal models suggests a greater proportion of caudate (vs. putamen) connectivity with visual cortex compared to auditory cortex (Seger 2013; Yeterian and Pandya 1995). Finally, to confirm these auditory corticostriatal findings in a larger dataset, we replicated our auditory corticostriatal analyses in participants from the Human Connectome Project with near-millimeter spatial resolution and high angular resolution (128 diffusion directions) collected in a single 7-Tesla MRI session.

To anticipate, complementary findings across datasets are consistent in delineating the structural pathways between human auditory cortex and dorsal striatum and suggests a privileged position for putamen in the larger auditory processing network. Over the last two decades, tremendous progress has been made in understanding the functional role of cortical pathways that map learned sounds (e.g., native speech) onto meaning (ventral pathway) and articulation/location (dorsal pathway). We posit that auditory-putamen streams are likely an important substrate underlying behaviorally-relevant auditory learning, and our results here provide a useful starting point for more extensive physiological examination.

## Methods

### Diffusion MRI data acquisition

We investigated auditory–striatal connectivity in two public diffusion MRI datasets. The first was collected in a single individual (male, approximately 30 years of age) over 18 hours of scanning on the MGH Connectome scanner. This machine has a 3 Tesla (3T) magnetic field but is optimized for diffusion MRI data acquisition, with high maximum gradient strength and slew rate enabling unparalleled diffusion MRI data quality in living human participants (Wang et al. 2021). 2808 volumes of diffusion MRI data were collected at 760 μm isotropic resolution, with b=0, 1000, and 2500 s/mm^2^. This data collection was approved by the Institutional Review Board of Partners Healthcare, and the participant provided written informed consent. Preprocessed data used in the present study are available from https://brain.labsolver.org/mgh_760.html (courtesy of F.-C. Yeh); data from the original authors are available at https://doi.org/10.5061/dryad.nzs7h44q2 (Wang et al. 2020).

To confirm our corticostriatal connectivity patterns across a larger sample, we included a second dataset comprised of 13 participants (8 female, 5 male, age range: 22–35 years of age) from the Human Connectome Project (HCP) Young Adult 7T dataset (Vu et al. 2015; Elam et al. 2021). We selected the lowest-numbered participant IDs in the 7T database without consideration of any demographic or other participant information. Data for each participant were acquired over approximately 1 hour of scanning at 1.05 mm isotropic resolution with a two-shell diffusion scheme (b=1000 and 2000 s/mm^2^) in 64 directions acquired twice at each shell, plus 15 b=0 volumes. Data collection for the HCP WU–Minn Consortium was approved by the institutional review boards of Washington University and the University of Minnesota, and participants provided written informed consent.

### Diffusion MRI processing

Preprocessed data from the sub-millimeter single-participant dataset was reconstructed in MNI space using q-space diffeomorphic reconstruction (QSDR) to obtain the spin distribution function, while the near-millimeter HCP 7T data were reconstructed in participant-native space with generalized q-sampling imaging (GQI) (Yeh et al. 2010; Yeh et al. 2021).

We then ran deterministic tractography using auditory cortical and striatal regions of interest (see below) as endpoints, generating 100,000 total tractography streamlines. For tractography, we defined the termination threshold at quantitative anisotropy = 0.01 (which quantifies streamline orientation-specific spin distribution functions), angular threshold = 45°, step size = 0.5, minimum streamline length = 5 mm, and maximum streamline length = 300 mm (Yeh et al. 2010; Yeh et al. 2021).

While deterministic tractography has demonstrated successful tracking between carefully defined regions of interest (Sarwar et al. 2019), particularly when using spin distribution function methods such as QSDR and GQI (Yeh 2022), probabilistic tractography can be more sensitive to ground truth connections across the brain (Grisot et al. 2021; Rosen and Halgren 2021; Farquharson et al. 2013). To assess the impact of tractography approach (deterministic or probabilistic), we further conducted Mrtrix3-based multi-shell, multi-tissue constrained spherical deconvolution with probabilistic tractography in the single-participant dataset in participant-native space. We generated 10 million probabilistic streamlines across the whole brain with Mrtrix3’s ‘tckgen’ command (tracking algorithm: iFOD2 [Second-order Integration over Fiber Orientation Distributions]; default parameters: angular threshold = 45°, step size = 0.5, minimum streamline length = 5 × voxel size, and maximum streamline length = 100 × voxel size.

### Connectivity analysis

We computed unthresholded connectivity matrices in DSI Studio (for the main deterministic tractography experiments) or Mrtrix3 (for the probabilistic tractography experiment) using the whole brain tractography streamlines and our predefined regions of interest as endpoints (see below). For all experiments, we then plotted connectivity heatmaps using Python’s matplotlib and seaborn packages (Hunter 2007; Waskom 2021).

### Region of interest segmentation

Striatal and cortical regions of interest were selected to mirror prior work examining regionspecific connections between dorsal striatum and auditory cortex in non-human primates (Yeterian and Pandya 1998). We extracted MNI-space subcortical segmentations from FreeSurfer (Dale et al. 1999) using dorsal striatal segmentations of the caudate and putamen.

Cortical segmentations of superior temporal cortex were extracted from the Human Connectome Project Multi-Modal Parcellation atlas (Glasser et al. 2016). The regions were from early auditory cortex: A1, lateral belt (LBelt), medial belt (MBelt), parabelt (PBelt), and retro-insular area (RI); auditory association cortex: A4, A5, dorsal anterior superior temporal sulcus (STSda), dorsal posterior superior temporal sulcus (STSdp), ventral anterior superior temporal sulcus (STSva), ventral posterior superior temporal sulcus (STSvp), anterior superior temporal gyrus (STGa), and anteromedial planum polare (TA2), and temporal pole (TGd). We also extracted visual regions of interest (V1, V2, V3, and V4) for a cross-modal comparison.

## Results

### Sub-millimeter single participant results

In the single-participant MGH 760 μm dataset, 100,000 streamlines were generated that had endpoints in the auditory cortical and striatal regions of interest (Figure 1). There were significantly more connections between superior temporal cortex and putamen than between superior temporal cortex and caudate (paired t-test: t = 4.061, p = 0.0004). Putamen streamline counts were higher than caudate streamline counts for all superior temporal cortical regions, regardless of the size of the region of interest. Caudate was connected most strongly with anterior superior temporal cortex and temporal pole as compared to primary and secondary auditory cortex more posteriorly (Figure 1C). Meanwhile, putamen was broadly connected with regions across superior temporal cortex, from primary auditory cortex regions posteriorly to associative regions more anteriorly. Streamlines to sulcal regions of superior temporal cortex tended to be more minimal than to gyral portions of the cortex.

**Figure 1.**
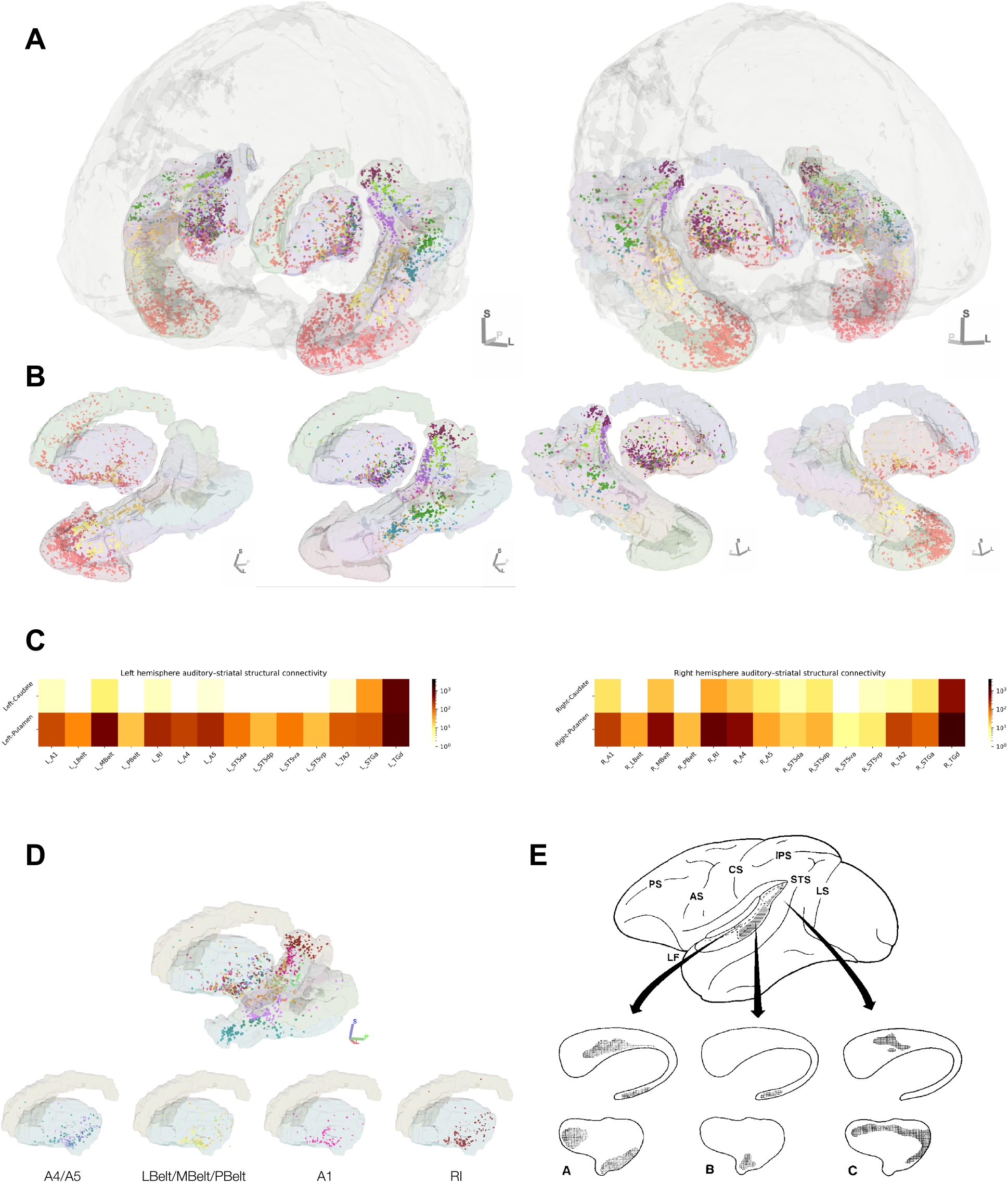
Tractography streamline endpoints in the sub-millimeter single-participant dataset. Endpoints are color-coded by cortical endpoint; for example, orange points denote streamlines connecting dorsal striatum and temporal pole (TGd), while maroon corresponds to retroinsular (RI) cortical connections. A: all corticostriatal endpoints, angled so that the left hemisphere (top left) and right hemisphere (top right) are visible. B: endpoints from anterior superior temporal cortex and temporal pole (bottom far left and bottom far right) and from posterior and medial superior temporal cortex (bottom second from left and second from right). C: streamline counts to caudate and putamen from each superior temporal cortical region (displayed in log scale). D: Tractography endpoints between dorsal striatum and posteromedial superior temporal cortex (auditory core and belt) in the submillimeter single-participant dataset. E: primary auditory cortical terminations in dorsal striatum, adapted from Yeterian and Pandya (1998).

### Striatal termination points of auditory cortical streamlines

We investigated the specific endpoints of auditory corticostriatal streamlines. When looking at striatal terminations, only anterior superior temporal cortex exhibited meaningful connections with caudate head and rostral putamen (Figure 1A). The other divisions of superior temporal cortex were strongly connected with caudal putamen but showed limited connectivity with rostral putamen or caudate.

Looking more focally at the auditory core and its surrounding structures, we visualized the dorsal striatal endpoints of streamlines that ended specifically in posteromedial superior temporal cortex (Figure 1D). Within putamen most streamlines ended in the caudal half, with many of these terminating on the ventral aspect of putamen. Minimal streamlines reached caudate.

### Probabilistic tractography

We next ran probabilistic tractography using a separate analysis pipeline to confirm that our auditory corticostriatal connectivity results hold across different analytical approaches. As validation of our findings using deterministic tractography, we found strongest probabilistic tractography connectivity patterns between putamen and superior temporal gyrus, with minimal connectivity between caudate and posterior STG or HG (Figure 2).

**Figure 2.**
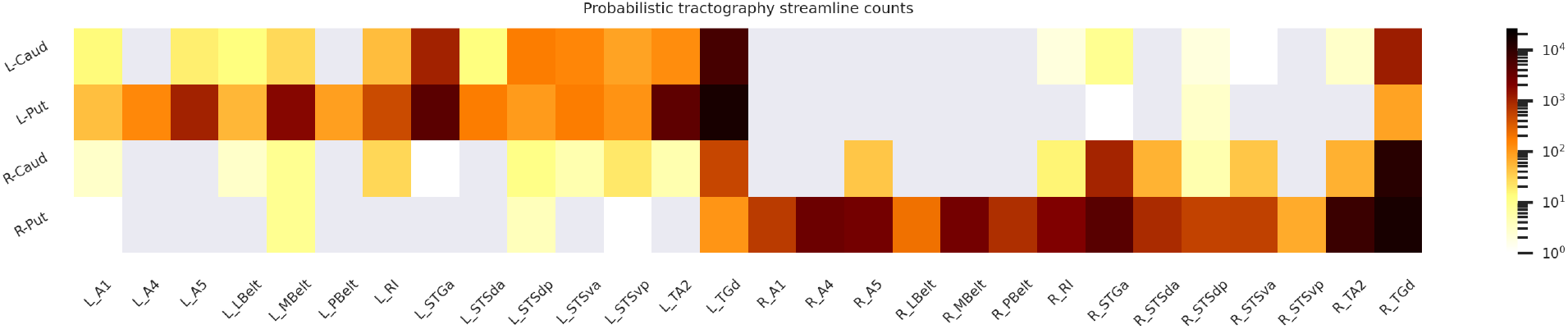
Probabilistic tractography streamline counts (displayed in log scale) between dorsal striatal and superior temporal cortical regions.

### Near-millimeter group results

Finally, we sought to replicate key findings in a separate high-quality dataset collected with 7T MRI as a part of the Human Connectome Project (HCP). Similar to the sub-millimeter connectivity results, when averaging across the near-millimeter HCP 7T group, auditory corticostriatal connectivity was much denser to putamen than to caudate (Figure 3, top row). In particular, caudate almost exclusively connected to anterior-most temporal regions. Again, superior temporal sulcus was less connected with putamen than other regions of superior temporal cortex.

**Figure 3.**
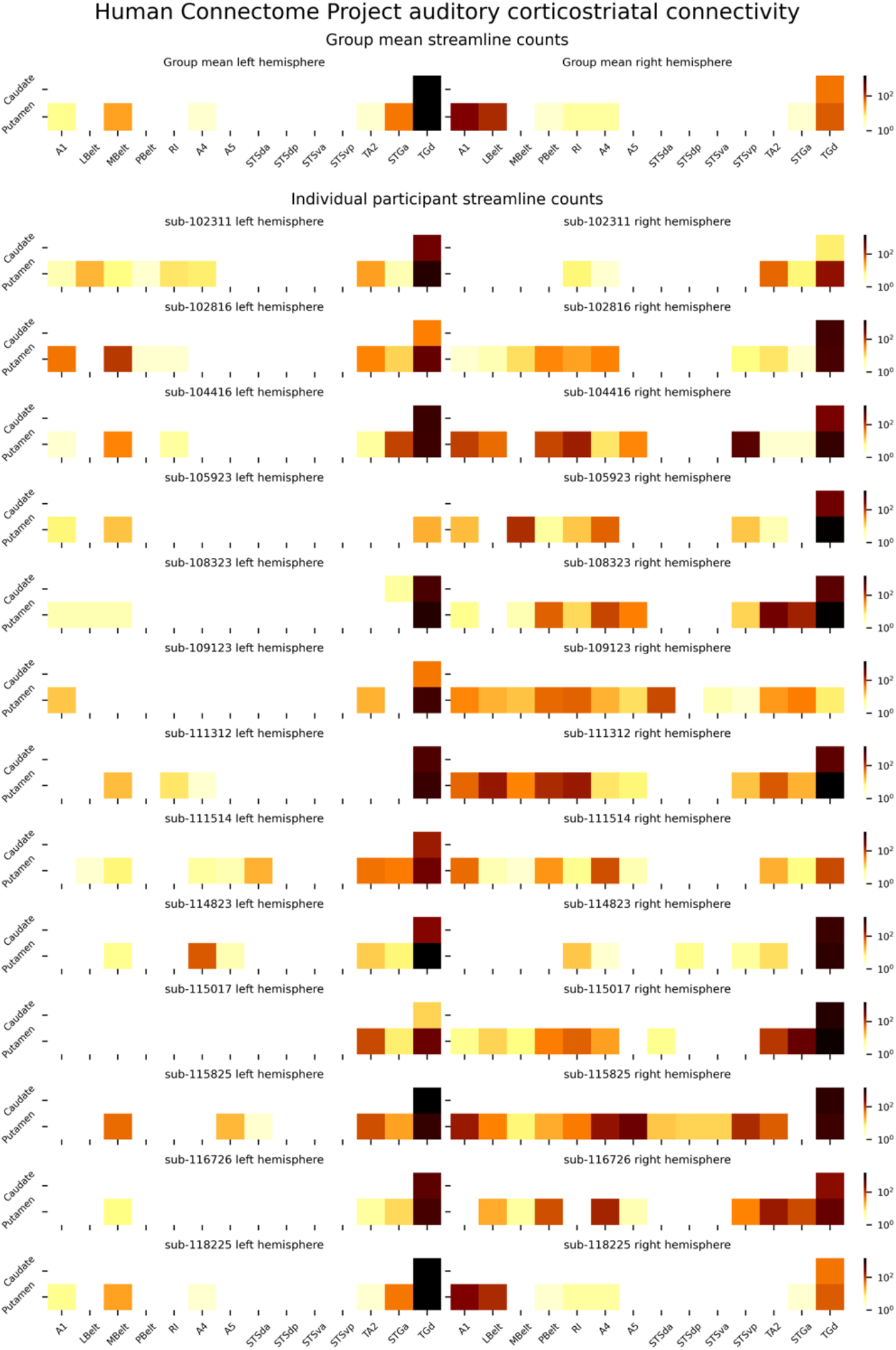
Top: Mean streamline connectivity across 13 participants from the Human Connectome Project 7T dataset. Bottom: Streamline count heatmaps for each individual included in the near-millimeter Human Connectome Project 7T dataset. Values are plotted on a log scale.

Across individuals, connectivity patterns were highly consistent (Fig. 5, remaining rows). Connectivity with putamen was higher across superior temporal cortex. Right hemisphere streamline counts were higher than left hemisphere counts across individuals.

### Visual corticostriatal results

Next we sought to assess the similarity of striatal projections across cortical sensory systems. We mapped corticostriatal connectivity in the visual system by generating tractography streamlines between major visual cortical regions and dorsal striatal structures (Figure 4). In comparison to auditory corticostriatal connectivity, visual corticostriatal streamline counts were greatly reduced. Additionally, unlike observations in putative auditory homologues, there were no significant differences between putamen and caudate visual corticostriatal streamline counts (paired t-test: t = 1.014, p = 0.3444). Visual corticostriatal streamlines reached more mixed striatal endpoints than auditory corticostriatal connections, with left V1–V3 reaching left striatal subdivisions fairly equally, while right V1–V3 had more limited connectivity with right putamen.

**Figure 4.**
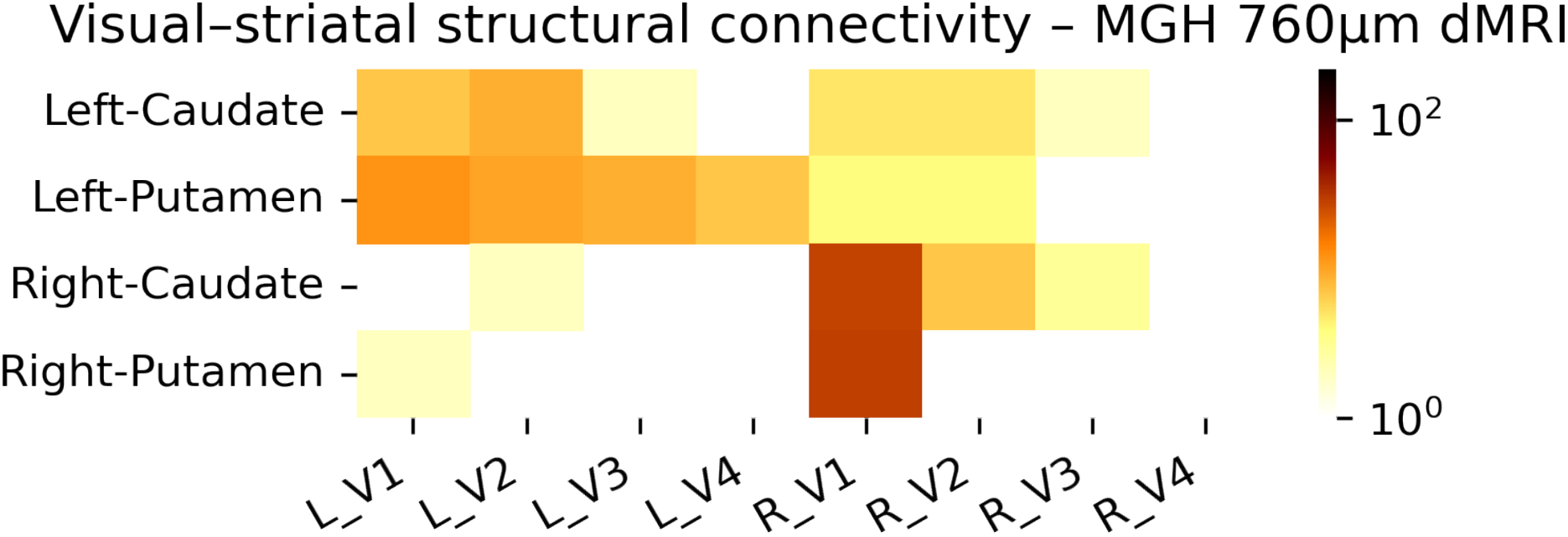
Streamline counts ending in striatal regions of interest and visual cortical regions of interest in the sub-millimeter single-participant dataset. Values are plotted on a log scale.

## Discussion

Building on animal evidence for the auditory corticostriatal pathway (Yeterian and Pandya 1998; Znamenskiy and Zador 2013), we used high resolution diffusion-weighted MRI tractography to estimate auditory corticostriatal connectivity in living humans. Our findings are broadly and specifically consistent with those of tract-tracing studies in non-human primates. In two distinct high-quality, high-resolution datasets, we found more widespread connectivity between putamen and superior temporal cortex than between caudate and superior temporal cortex. This finding was consistent across tractography methods (deterministic and probabilistic tractography). Auditory corticostriatal connectivity patterns differed from the pattern of visual corticostriatal connectivity, which did not mirror significant differences between caudate and putamen connectivity. The dramatic differences between caudate and putamen connectivity in the auditory system—and the lack of differences in the visual system—are unlikely to be due to differences in striatal volume, with caudate and putamen having comparable volumes (Abedelahi et al. 2013; Hokama et al. 1995). Further, specific subdivisions of dorsal striatum— putamen and head of the caudate—had divergent connectivity patterns with superior temporal cortex, with caudate head mainly connecting to anterior superior temporal cortex, and posterior auditory cortical regions preferentially connecting to caudal and ventral putamen. These findings may have implications in communication disorders associated with striatal structure and function (discussed more below). The general medial-lateral organization of corticostriatal topography in the present data reinforce the concept of “functional mosaicism” described by Selemon and Goldman-Rakic (Selemon and Goldman-Rakic 1985). Overall, the consistency of our connectivity findings across hemispheres, datasets, tractography methods, and individuals suggests that highquality, high-resolution diffusion MRI tractography is sensitive to fine-grained auditory corticostriatal connectivity patterns.

Diffusion MRI tractography is an inferential method that does not directly measure white matter. To our knowledge, there are no reports in the literature of temporal corticostriatal white matter dissections in the human brain to which we can compare the present findings. However, our assessment of structural connectivity between superior temporal cortex and dorsal striatum closely matches tract tracing results in macaque (Yeterian and Pandya 1998) and squirrel monkey (Borgmann and Jürgens 1999). Generally, our findings reaffirm that striatal connections are more plentiful with auditory association cortical regions as compared to primary auditory cortex.

Further, we found that the auditory core and belt regions were more strongly connected with putamen than with caudate. Indeed, while auditory core and belt exhibited streamlines to caudal and ventral putamen in our results, they showed almost no connectivity with caudate. Yeterian and Pandya (Yeterian and Pandya 1998) did observe limited auditory core and belt connections to caudate in macaque, but only to the medial caudate and caudal-most tail, respectively. The tail of the caudate is challenging to segment in human T1-weighted MR images and is largely absent from standard brain atlases, including the FreeSurfer segmentation used in this study (Fischl et al. 2002), which may explain some of the lacking auditory cortical–caudate streamlines. In light of this methodological limitation, it is possible that the sparse caudate connections present in macaques are in fact also present in humans (beyond the limited but present connections between auditory belt and body of the caudate in single-participant sub-millimeter dataset). In animal models, the caudal-most “tail of the striatum”—encompassing both the caudate tail and posterior putamen—has been hypothesized as a functionally distinct unit receiving multisensory inputs (Valjent and Gangarossa 2021; Cox and Witten 2019), which aligns with the Yeterian and Pandya results (Yeterian and Pandya 1998) and suggests that finer spatial resolution and more accurate caudate tail segmentation with human MRI could reveal similar tractography connections between the auditory core and tail of the caudate.

When looking more broadly across superior temporal cortex, Yeterian and Pandya (Yeterian and Pandya 1998) described heavy connections between anterior superior temporal lobe and anterior dorsal striatum (head of the caudate and rostral putamen). We found similar patterns in the single-participant sub-millimeter tractography results—specifically from temporal pole and anterior superior temporal gyrus to anterior caudate and putamen. Intriguingly, this anterior– posterior connectivity scheme is consistent with and could reasonably map onto known “what” and “where” pathways in the primate auditory cortex (Romanski et al. 1999; Rauschecker and Tian 2000; Rauschecker and Scott 2009; Kaas and Hackett 2000). Thus, corticostriatal connections may contribute to “cortical” functional networks such the dual auditory streams.

One notable difference between our current work and the previous tract-tracing work is that in the macaque, middle and posterior superior temporal cortex exhibited reduced but present caudate connections, which were largely—but not wholly—absent in both of our datasets. Thus, while our diffusion MRI tractography estimates of auditory corticostriatal connectivity broadly align with tract-tracing work in non-human primates (Yeterian and Pandya 1998), without the availability of ground-truth connectivity methods in humans, we cannot be certain whether the differences in our results are due to methodological limitations in human diffusion MRI tractography or due to actual evolutionary distinctions between primate species. Indeed, a recent diffusion MRI tractography comparing human and chimpanzee cortical language networks suggests an expansion of posterior temporal connectivity in humans, while anterior temporal connectivity is similar between species but with some expanded cortical targets such as through the uncinate fasciculus (Sierpowska et al. 2022). It is possible that corticostriatal connectivity would similarly retain core connectivity features over the course of primate evolution while also developing human-specific connections that support human auditory learning and decisionmaking.

In this work, we mapped structural connectivity between human superior temporal cortex and dorsal striatum using diffusion MRI tractography. While tractography can provide us with estimates of white matter *orientation*, it cannot tell us the *direction* of a given white matter tract. However, previous neuroanatomical investigations into corticostriatal systems in nonhuman primates provide evidence for multiple closed loops between cerebral cortex and dorsal striatum (Middleton and Strick 2000). While dorsal striatum receives input from across the cortex (Selemon and Goldman-Rakic 1985), in the temporal lobe, only inferotemporal cortex has been established as a recipient of basal ganglia outputs by way of substantia nigra pars reticulata (Middleton and Strick 1996). Thus, diffusion MRI tractography connections between superior temporal cortex and dorsal striatum in the present study are likely to represent corticostriatal white matter pathways as opposed to striatocortical pathways.

While literature in non-human primates and other animal models is highly suggestive of a central role for posterior putamen in auditory processing (Clarey and Irvine 1986; Zhong et al. 2014), in humans, the literature on auditory corticostriatal connectivity is limited. One study that focused on auditory learning using fMRI also estimated auditory corticostriatal connectivity using diffusion MRI (Feng, Yi, & Chandrasekaran, 2019). By using functional activation clusters in anterior STG and putamen as probabilistic tractography seeds, the investigators found strong connectivity between the two regions implicated in novel speech sound category learning.

Additionally, auditory corticostriatal connectivity is implicated in disorders that impact communication. With resting state functional MRI approaches, Gordon et al. (Gordon et al. 2021) mapped cortical functional connectivity with striatum, finding a putamen cluster that strongly connects functionally to the cortical language network. Hinkley et al. (Hinkley et al. 2015; Hinkley et al. 2021) similarly used functional connectivity to investigate corticostriatal interactions and found *ventral* striatal connectivity effects in tinnitus participants. A follow up study (Hinkley et al. 2021) found that targeted electrical stimulation of the caudal-most caudate tail provided significant benefits to participants with tinnitus.

In children with developmental stuttering, a disorder with both motor and perceptual deficits, decreased functional and structural connectivity with motor, speech, and auditory cortical regions has been observed (Chang and Zhu 2013). In adults who stutter, putamen exhibited higher quantitative MRI values that are associated with iron levels and could reflect elevated neurotransmitter levels in sensorimotor striatum (Cler et al. 2021). Developmental language disorders and dyslexia may similarly be impacted by abnormal striatal function during auditory behavior (Krishnan et al. 2016), with abnormal myelin-related quantitative MRI values in caudate specifically (Krishnan et al. 2022). However, despite broad consensus that dorsal striatal dysfunction can contribute to auditory deficits in humans, the precise pathways between auditory cortex and dorsal striatum that underlie such auditory function have not, until now, been clearly delineated. Aligning with the dual auditory streams hypothesis, our results suggest that dysfunction in anterior dorsal striatum may compromise higher-order auditory and language function via the anterior auditory corticostriatal pathways, while dysfunction in posterior dorsal striatum (particularly putamen) would more likely affect sensorimotor aspects of audition and communication via the posterior auditory corticostriatal pathways.

Across two high quality, high resolution human diffusion MRI datasets, the present results suggest a privileged position for putamen in auditory processing and learning. Indeed, previous functional research in humans identified a key role for putamen in learning new sound categories, whether they be speech or non-speech stimuli (Lim et al. 2014; Yi et al. 2016; Feng et al. 2019; Lim et al. 2019). Despite a lack of ground truth human connectivity and functional mapping, as well as some evidence that macaque and human striatum are not fully homologous (Liu et al. 2021), the non-invasive human imaging literature aligns well with research in animal models. Our own virtual dissection results closely follow tract-tracing work in non-human primates that distinguished anterior from posterior superior temporal connectivity with dorsal striatum (Yeterian and Pandya 1998). Future work will map specific auditory functions within human dorsal striatum, allowing us to probe the relationship between precise auditory corticostriatal connections and communication-related disorders such as autism spectrum disorders, developmental stuttering, and tinnitus.

## Acknowledgements

We thank Fuyixue Wang and coauthors for making their sub-millimeter single-participant diffusion MRI data publicly available, and Fang-Cheng Yeh for releasing the data in DSI Studio. Human Connectome Project data were provided [in part] by the Human Connectome Project, WU-Minn Consortium (Principal Investigators: David Van Essen and Kamil Ugurbil; 1U54MH091657) funded by the 16NIH Institutes and Centers that support the NIH Blueprint for Neuroscience Research; and by theMcDonnell Center for Systems Neuroscience at Washington University. Funding was provided by NIHNIDCD awards K01DC019421 (KRS) and R01DC015504 (BC and KRS).

## Statements and Declarations

### Author contributions

All authors contributed to the study conception and design. Data analysis was performed by KRS. The first draft of the manuscript was written by KRS and all authors commented on previous versions of the manuscript. All authors read and approved the final manuscript.

### Data availability

The data used in this study are available publicly. For the sub-millimeter dataset, preprocessed data are available from https://brain.labsolver.org/mgh_760.html; data from the original authors are available at https://doi.org/10.5061/dryad.nzs7h44q2. Human Connectome Project data are available from https://db.humanconnectome.org.

### Ethics approval

This study was performed in line with the principles of the Declaration of Helsinki. For the sub-millimeter dataset, data collection was approved by the Institutional Review Board of Partners Healthcare, and the participant provided written informed consent. Data collection for the Human Connectome Project WU–Minn Consortium was approved by the institutional review boards of Washington University and the University of Minnesota, and participants provided written informed consent.

### Consent to participate

Informed consent was obtained from all individual participants included in the study.

